# sxLaep: a Lightweight and Accurate Enzyme Predictor for High-throughput Mining of Metagenomic Sequences

**DOI:** 10.64898/2026.05.06.723393

**Authors:** Hongyu Duan, Xinyu Han, Yijun Mo, Bozhen Ren, Li C. Xia

## Abstract

**Motivation:** Metagenomic sequencing generates petabyte-scale sequence datasets that strain both deep learning and alignment based enzyme annotation tools. A lightweight rapid and accurate filter tool is needed to filter and identify enzymatic sequences prior to resource-intensive functional prediction.

**Results:** We present sxLaep (Lightweight and Accurate Enzyme Predictor), a resource-efficient framework using lightweight physicochemical features for enzyme pre-screening. On the external validation set, sxLaep completed prediction in only 0.002 s/sequence, which is 22.9-fold faster than Diamond (0.0457 s/sequence). It used 372.16 MB peak memory, corresponding to a 54.4% memory reduction relative to Diamond (815.64 MB). sxLaep achieved an accuracy of 99.34% and the highest recall in remote homology detection, including enzyme candidates missed by alignment-based methods. We further successfully applied sxLaep to a marine metagenomic enzyme-mining workflow, demonstrating its utility for high-throughput discovery from large-scale metagenomic sequences.

**Availability and Implementation:** sxLaep is available as a Python package at https://pypi.org/project/sxlaep and is maintained as an open-source software repository at https://github.com/labxscut/sxLaep. Detailed installation, usage, and Docker deployment instructions are provided in the GitHub repository to support reproducible enzyme prediction and model execution.

## Introduction

Enzymes are crucial building blocks of bio-design and biomanufacturing. Massive metagenomic sequencing projects have enabled exploration of previously inaccessible novel enzymes with considerable industrial and therapeutic potential (Danko *et al*., 2021, Thompson et al., 2017, Muammar et al., 2026, Wani et al., 2026, Kumar et al., 2025, Alam et al., 2025). For example, targeted mining workflows have enabled large-scale metagenomic enzyme discovery from the oceans (Xu *et al*., 2025, Moorhoff *et al*., 2026) with many downstream applications.

However, the exponential growth of metagenomic datasets, while expected to accelerate with third-generation sequencing platforms (Sereika *et al*., 2025, Sun et al., 2026, Benoit et al., 2026), already poses significant pre-screening challenges to existing enzyme function predictors. Alignment-based tools such as BLAST (Altschul *et al*., 1990, Al-Fatlawi et al., 2023) and Diamond (Buchfink *et al*., 2021) are computationally costly at metagenomic scale and demonstrate low sensitivity in detecting remote homologs, limiting their utility in unexplored microbial environments, where functional diversity is substantial and available homologous references are limited (Pavlopoulos *et al*., 2023).

Recent deep learning approaches, including large protein language models for sequence representation and structure inference (Lin *et al*., 2023), contrastive learning for enzyme function such as CLEAN (Yu *et al*., 2023), and structure-aware function annotation (Derry *et al*., 2025), achieve strong accuracy on benchmarks. However, neural encoding and structure prediction remain memory- and compute-intensive (Jumper *et al*., 2021), making exhaustive pre-screening of petabyte-scale metagenomic data impractical on typical infrastructure.

This gap motivates lightweight pre-screening tools that rapidly distinguish enzymatic from non-enzymatic sequences and reduce computational overhead in metagenomic mining workflows (Robinson *et al*., 2021). We introduce sxLaep (Lightweight and Accurate Enzyme Predictor), a specialized pre-screener that balances efficiency and accuracy using physicochemical descriptors: pseudo-amino-acid composition (PseAAC) (Huang and Zhang, 2024), composition-transition-distribution (CTD) (Zulfiqar *et al*., 2021), and windowed amino-acid composition (WinAAC) (Vedula *et al*., 2025), in place of high-dimensional neural embeddings. Coupled to an optimized gradient-boosting classifier, this “filter-then-validate” design supports rapid screening of very large sequence sets while retaining candidates with weak primary-sequence similarity. We benchmark sxLaep against alignment and deep-learning baselines and apply it at scale to marine metagenome-assembled genomes in line with global marine microbial diversity and bioprospecting efforts (Chen *et al*., 2024).

## Implementation

### Overview

We present sxLaep, a fast, accurate, and lightweight framework for large-scale metagenomic analysis that combines streamlined feature extraction with an optimized gradient boosting classifier while avoiding GPU-intensive embeddings and preserving remote-homology detection capability.

### Feature Extraction Strategy

sxLaep maps variable-length protein sequences to fixed-length feature vectors by integrating three complementary feature groups that capture both global and local physicochemical patterns of real enzymes. For clarity, these feature groups can be organized into two categories: (i) a **global** descriptor set (PseAAC) and (ii) a **local** descriptor set (CTD and WinAAC (see Supplementary Note S5)).

#### Global Descriptors: Pseudo-Amino Acid Composition (PseAAC)

sxLaep incorporates PseAAC, which are sequence-order correlation factors derived from physicochemical properties (hydrophobicity, hydrophilicity, side-chain mass). For a protein sequence *P* = *R*_1_*R*_2_ … *R*_*L*_, the PseAAC feature vector is:

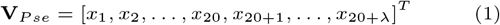

where the components are:

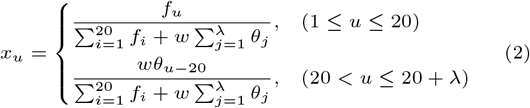

Here, *f*_*u*_ is the normalized frequency of the *u*-th amino acid, *w* = 0.05 balances composition and order effects, and 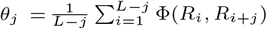 measures physicochemical distances. This ensures global structural tendencies are preserved even with low sequence identity.

#### Local Pattern Descriptors: Composition, Transition, and Distribution (CTD)

sxLaep also incorporates CTD, which categorizes amino acids into three groups (polar, neutral, hydrophobic) and calculates three descriptors:

- **Composition (C):** Global percentage of each group:

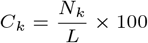
- **Transition (T):** Frequency of group changes:

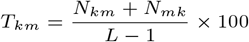
- **Distribution (D):** Spatial profile along the backbone:

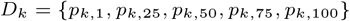

These descriptors capture catalytic triads, hydrophobic cores, and other local enzymatic motifs.

### Model Architecture and Optimization

sxLaep employs XGBoost as core classifier, selected for its scalability and ability to handle large sparse data (see Supplementary Figure S1). To minimize false negatives (more detrimental than false positives for pre-screening), we formulated a customized objective function, where the decision threshold *τ* was tuned via cross-validated grid search:

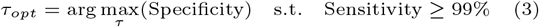

This ensures FNR *<* 1%, guaranteeing retention of potential enzymes with low sequence identity for downstream analysis.

## Results

### Robust Identification of Remote Homologs

A key challenge in metagenomics-based enzyme discovery is identifying the large pool of functionally uncharacterized microbial proteins, often referred to as functional “dark matter” (Pavlopoulos *et al*., 2023, Zhang *et al*., 2025, Xian *et al*., 2025). This broader hidden functional space has also been emphasized in recent dark-matter studies beyond cellular metagenomes, including viral sequence space (Kosmopoulos and Anantharaman, 2025). To evaluate sxLaep’s capability in this task, we stratified the test set based on sequence identity relative to the training data. As illustrated in Fig. 1b, we compared the recall (sensitivity) of sxLaep against Diamond (BLASTp-compatible). In high-identity regions (*>* 40%), both methods demonstrated comparable performance with near-perfect recall. In contrast, sxLaep significantly outperforms Diamond in the low-identity region (*<* 20% sequence identity). In this remote homology region (0–20% identity), Diamond recall dropped to 0, whereas sxLaep maintained a recall of 0.471. This performance gap confirms that sxLaep successfully captures conserved physicochemical patterns (e.g., hydrophobicity profiles and charge distributions) that persist even when primary sequence identity vanishes, making it significantly more effective for mining novel biocatalysts.

**Fig. 1.**
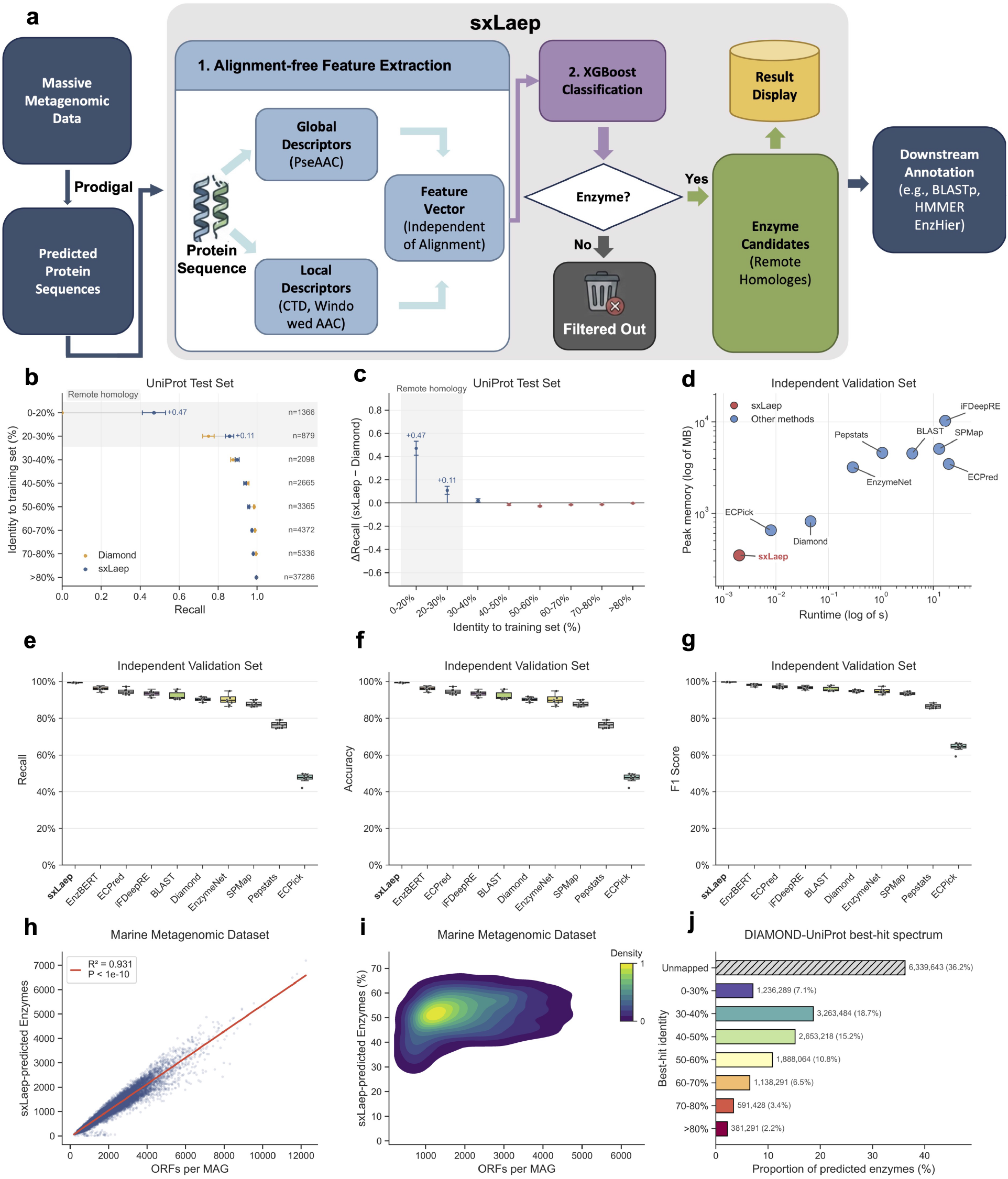
Overview of the sxLaep pre-screening workflow, benchmark performance, and ecological patterns across large-scale marine MAG analyses: (a) the sxLaep pre-screening workflow; (b) recall across sequence-identity bins in the UniProt test set; (c) ∆ Recall relative to Diamond (sxLaep - Diamond); (d) runtime versus peak memory usage on the independent validation set (log scale); (e) recall on the independent validation set; (f) accuracy on the independent validation set; (g) F1-score on the independent validation set; (h) relationship between ORFs per MAG and enzyme-coding ORFs; (i) bivariate density of ORFs per MAG and proportion of enzyme-coding ORFs; (j) DIAMOND-UniProt best-hit spectrum of predicted enzyme-coding ORFs.

### High Computational Efficiency and Scalability

We benchmarked sxLaep against state-of-the-art tools, including alignment-based methods (BLAST (Altschul *et al*., 1990), Diamond (Buchfink *et al*., 2021)), classical sequence-statistics baselines (Peptstats (Dalkiran *et al*., 2018)), and representative learning-based baselines (iFDeepRE (Tan *et al*., 2024), EnzymeNet (Watanabe *et al*., 2023), ECPICK (Han *et al*., 2024), ECPred (Dalkiran *et al*., 2018), and SPMap (Dalkiran *et al*., 2018)). As shown in Fig. 1d, sxLaep achieves minimal resource requirements compared to competing approaches. sxLaep required 0.002 s/sequence, making it 4-fold faster than the second-fastest learning-based baseline ECPICK (0.008 s/sequence), 1,981-fold faster than BLAST (3.962 s/sequence), and 22.9-fold faster than Diamond (0.0457 s/sequence). sxLaep also used only 372.16 MB peak memory, corresponding to a 42.9% reduction relative to ECPICK (651.76 MB), a 91.8% reduction relative to BLAST (4523.01 MB), and a 54.4% reduction relative to Diamond (815.64 MB). This resource efficiency enables high-throughput screening on standard computing hardware without incurring computational bottlenecks in metagenomic workflows.

### High Sensitivity, Accuracy, and Reliability

High recall is critical for pre-screening, as missing potential candidates is more costly than admitting extra false positives for downstream validation. Fig. 1e shows that sxLaep achieved a mean recall of 99.4%, compared with 96.1% for the runner-up EnzBERT (Buton *et al*., 2023), representing a 3.3-percentage-point improvement. This performance ensures retention of enzymatic candidates with weak sequence identity for functional analysis.

Beyond its superior recall, sxLaep maintained a mean accuracy of 99.34% and a mean F1-score of 99.7% (Figs. 1f-g). In the Diamond-inclusive comparison benchmark, sxLaep also outperformed Diamond, improving mean recall from 90.2% to 99.4%, mean accuracy from 90.2% to 99.34%, and F1-score from 94.9% to 99.7%. Together with the remote-homology advantage in Fig. 1b, these results indicate that sxLaep preserves strong global classification performance while remaining substantially more sensitive in low-identity sequence space.

### Enzyme Distribution in Marine MAGs

To examine the broader applicability of sxLaep, we analyzed predicted enzymes across 16,240 marine MAGs (Chen *et al*., 2024). As shown in Figs. 1h-i, sxLaep predicted MAG-wise enzyme counts scale linearly with MAG (ORF) sizes, consistent with the well-known proportional relationship between genome-scale gene content and metabolic capacity as represented by enzyme counts (Konstantinidis and Tiedje, 2004). Fig. 1j further summarizes the DIAMOND-UniProt best-hit spectrum of predicted enzyme-coding ORFs. Using DIAMOND searches with an e-value cutoff of 1e-3 and retaining one best hit per query by lowest e-value, highest bitscore, and highest sequence identity, we found that 36.2% of predicted enzymes had no retained UniProt hit, while mapped sequences were concentrated in low-to moderate-identity bins. This pattern suggests that a substantial fraction of sxLaep-predicted enzymes correspond to database-remote or currently UniProt-unmapped candidates. Details of the analyses are provided in the Supplementary Note S1.

## Conclusion

sxLaep demonstrates significant computational efficiency (22.9-fold faster and 54.4% lower peak memory than Diamond) while achieving 99.34% accuracy and the best recall in remote homology detection—addressing a computational bottleneck in metagenomics-based enzyme discovery where rapid pre-screening is essential prior to functional validation. With metagenomic datasets expanding with advancing sequencing technologies, sxLaep’s lightweight architecture enables researchers to rapidly pre-screen billions of proteins on standard hardware, identifying enzymatic candidates for downstream functional assays and heterologous expression pipelines. This “filter-then-validate” tool is well-suited for industrial biocatalyst discovery and large-scale functional genomics projects where functional assays are resource-intensive. We expect sxLaep to be a practical utility in metagenomic annotation workflows, particularly valuable where remote homologs are diverse.

## Acknowledgements

This study was funded by the National Science Foundation of China (12571529) and Guangdong Basic and Applied Basic Research Foundation (2024A1515-010699) to LCX.

