## Supplementary Figure S1 for "sxLaep: a Lightweight and Accurate Enzyme Predictor for High-throughput Mining of Metagenomic Sequences"

#### Supplementary Figures

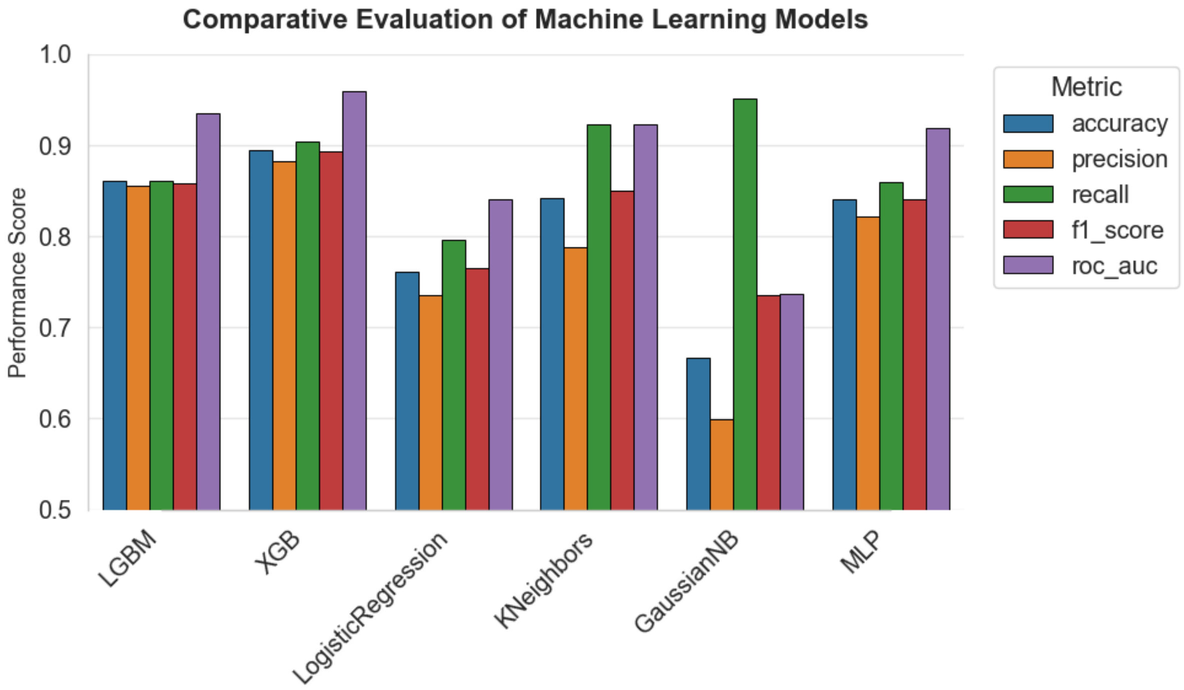


**Figure S1. Comparative evaluation of machine learning models.** Figure S1 compares six conventional machine learning models using accuracy, precision, recall, F1-score, and ROC-AUC.

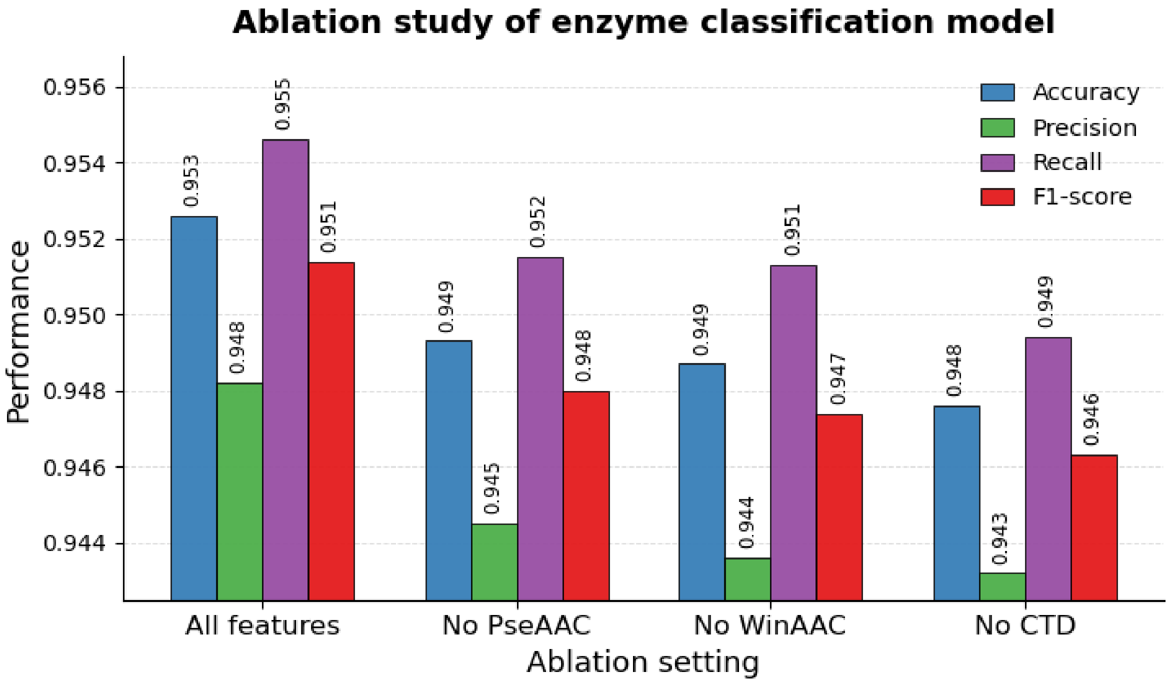


**Figure S2. Ablation study of the sxLaep feature design.** Figure S2 shows the contribution of the three feature groups used in sxLaep. Removing any single component reduced performance relative to the full model, confirming that PseAAC, WinAAC, and CTD all contributed complementary signal.

### Supplementary Tables

**Table S1. Comparative evaluation of machine learning models.**

| Model | Accuracy | Precision | Recall | F1-score | ROC-AUC |
|---|---|---|---|---|---|
| LGBM | 0.8613 | 0.8548 | 0.8609 | 0.8578 | 0.9352 |
| XGBoost | 0.8944 | 0.8819 | 0.9036 | 0.8927 | 0.9593 |
| LogisticRegression | 0.7616 | 0.7353 | 0.7960 | 0.7644 | 0.8400 |
| KNeighbors | 0.8415 | 0.7874 | 0.9232 | 0.8499 | 0.9234 |
| GaussianNB | 0.6668 | 0.5989 | 0.9516 | 0.7352 | 0.7366 |
| MLP | 0.8410 | 0.8213 | 0.8600 | 0.8402 | 0.9192 |

**Table S2. Ablation study of the sxLaep feature design.**

| Ablation setting | Accuracy | Precision | Recall | F1-score |
|---|---|---|---|---|
| All features | **0.9526** | **0.9482** | **0.9546** | **0.9514** |
| No PseAAC | 0.9493 | 0.9445 | 0.9515 | 0.9480 |
| No WinAAC | 0.9487 | 0.9436 | 95.13% | 0.9474 |
| No CTD | 0.9476 | 0.9432 | 0.9494 | 0.9463 |

**Table S3. Remote-homology recall by sequence identity bin (Part 1 of 2).**

| Identity bin | Total sequences | Positive sequences | sxLaep recall | sxLaep SE |
|---|---|---|---|---|
| 0–20% | 1366 | 68 | **0.4706** | 0.0599 |
| 20–30% | 879 | 240 | **0.8583** | 0.0217 |
| 30–40% | 2098 | 784 | 0.8980 | 0.0106 |
| 40–50% | 2665 | 1226 | 0.9380 | 0.0069 |
| 50–60% | 3365 | 1751 | 0.9572 | 0.0048 |
| 60–70% | 4372 | 2404 | 0.9738 | 0.0032 |
| 70–80% | 5336 | 2817 | 0.9808 | 0.0026 |
| > 80% | 37286 | 18591 | 0.9960 | 0.0005 |

**Table S3. Remote-homology recall by sequence identity bin (Part 2 of 2).**

| Identity bin | Diamond recall | Diamond SE | Delta recall (sxLaep - Diamond) | Delta SE |
|---|---|---|---|---|
| 0–20% | 0.0000 | 0.0000 | **0.4706** | 0.0599 |
| 20–30% | 0.7500 | 0.0291 | **0.1083** | 0.0341 |
| 30–40% | 0.8763 | 0.0119 | 0.0217 | 0.0147 |
| 40–50% | 0.9519 | 0.0061 | -0.0139 | 0.0080 |
| 50–60% | 0.9852 | 0.0030 | -0.0280 | 0.0052 |
| 60–70% | 0.9896 | 0.0020 | -0.0158 | 0.0037 |
| 70–80% | 0.9947 | 0.0013 | -0.0138 | 0.0026 |
| > 80% | 0.9982 | 0.0003 | -0.0023 | 0.0005 |

**Table S4. Classification performance summary (Part 1 of 2).**

| Model | Mean FNR | FNR SD | Accuracy mean | Accuracy SD |
|---|---|---|---|---|
| sxLaep | **0.0064** | 0.0045 | **99.34%** | 0.0045 |
| EnzBERT | 0.0388 | 0.0110 | 0.9612 | 0.0110 |
| ECPred | 0.0551 | 0.0135 | 0.9449 | 0.0135 |
| iFDeepRE | 0.0648 | 0.0143 | 0.9352 | 0.0143 |
| BLAST | 0.0786 | 0.0215 | 0.9214 | 0.0215 |
| Diamond | 0.0977 | 0.0103 | 0.9023 | 0.0103 |
| EnzymeNet | 0.0992 | 0.0250 | 0.9008 | 0.0250 |
| SPMap | 0.1217 | 0.0135 | 0.8783 | 0.0135 |
| Pepstats | 0.2357 | 0.0171 | 0.7643 | 0.0171 |
| ECPick | 0.5255 | 0.0228 | 0.4745 | 0.0228 |

**Table S4. Classification performance summary (Part 2 of 2).**

| Model | Recall mean | Recall SD | F1 mean | F1 SD |
|---|---|---|---|---|
| sxLaep | **0.9936** | 0.0045 | **0.9968** | 0.0023 |
| EnzBERT | 0.9612 | 0.0110 | 0.9802 | 0.0057 |
| ECPred | 0.9449 | 0.0135 | 0.9716 | 0.0071 |
| iFDeepRE | 0.9352 | 0.0143 | 0.9665 | 0.0076 |
| BLAST | 0.9214 | 0.0215 | 0.9590 | 0.0115 |
| Diamond | 0.9023 | 0.0103 | 0.9486 | 0.0057 |
| EnzymeNet | 0.9008 | 0.0250 | 0.9476 | 0.0138 |
| SPMap | 0.8783 | 0.0135 | 0.9352 | 0.0076 |
| Pepstats | 0.7643 | 0.0171 | 0.8663 | 0.0110 |
| ECPick | 0.4745 | 0.0228 | 0.6433 | 0.0213 |

**Table S5. Runtime and peak memory usage.**

| Method | Runtime (s) | Peak memory (MB) |
|---|---|---|
| sxLaep | **0.002** | **372.16** |
| ECPick | 0.008 | 651.76 |
| EnzymeNet | 0.292 | 3192.30 |
| Pepstats | 1.066 | 4598.78 |
| BLAST | 3.962 | 4523.01 |
| SPMap | 12.907 | 5080.06 |
| iFDeepRE | 16.820 | 10290.75 |
| Diamond | 0.0457 | 815.64 |
| ECPred | 49.500 | 3763.00 |

**Table S6. Feature dimension overview.**

| Feature type | Dimensions | Description |
|---|---|---|
| PseAAC | 51 | Amino acid composition + sequence-order correlation + sequence length |
| CTD | 147 | Composition, transition, and distribution descriptors across 7 physicochemical properties |
| WinAAC | 60 | Segment-wise amino acid composition across 3 sequence windows |
| **Total** | **258** | **Integrated input feature space used by sxLaep** |

**Table S7. Amino acid grouping schemes used by CTD features.**

| Property | Group 1 | Group 2 | Group 3 |
|---|---|---|---|
| Hydrophobicity | RKEDQN | GASTPHY | CLVIMFW |
| Normalized van der Waals volume | GASTPD | NVEQIL | MHKFRYW |
| Polarizability | GASDT | CPNVEQIL | KMHFRYW |
| Secondary structure propensity | NDEQST | AILMV | CFGHKPRYW |
| Solvent accessibility | ALFCGIVW | RKQEND | MPSTHY |
| Polarity | LIFWCMVY | PATGS ANCQGHILMFPSTW | HQRKNED |
| Charge | KR | YV | DE |

### Supplementary Notes

#### Note S1. Enzyme Distribution in Marine MAGs

Here we provide additional detailed interpretation of the analysis results summarized in the main text for 16,240 marine MAGs (Chen et al., 2024), to further support the broader applicability of sxLaep. In this analysis, total ORF count is used as a practical proxy for genome-scale gene content.

As shown in Fig. 1h of the main text, predicted enzyme counts scale linearly with MAG (ORF) sizes, consistent with the well-known proportional relationship between genome-scale gene content and metabolic capacity as represented by enzyme counts (Konstantinidis and Tiedje, 2004). Fig. 1i further identifies a dominant density core centered around MAGs with approximately 2,000 total ORFs and an enzyme fraction near 52%, suggesting a common ecological configuration in the sampled marine genomes that is consistent with genomic niche differentiation (Rocap et al., 2003).

Fig. 1j shows the DIAMOND-UniProt best-hit spectrum of predicted enzyme-coding ORFs. DIAMOND searches were performed with an e-value cutoff of 1e-3. For each query, only one best hit was retained, prioritized by the lowest e-value, highest bitscore, and highest sequence identity. Bitscore was used only for ranking, not as an independent cutoff. Sequence identity was grouped into 0–30%, 30–40%, 40–50%, 50–60%, 60–70%, 70–80%, and > 80% bins. Overall, 36.2% of predicted enzymes had no retained UniProt hit, and the mapped sequences were concentrated in low- to moderate-identity bins, indicating a large fraction of database-remote or UniProt-unmapped enzyme candidates.

#### Note S2. Feature representation summary

sxLaep uses a 258-dimensional handcrafted feature representation that integrates global sequence composition, physicochemical correlation, and local compositional patterns. The full feature space is composed of three groups:

- `PseAAC` (51 dimensions): amino acid composition, sequence-order correlation, and sequence length.
- `CTD` (147 dimensions): composition, transition, and distribution descriptors across seven physicochemical grouping schemes.
- `WinAAC` (60 dimensions): amino acid composition computed over three sequence segments.

#### Note S3. Detailed definition of PseAAC features

The `PseAAC` block contains 51 features:

- 20 amino acid composition (`AAC`) features representing the relative frequencies of the standard amino acids.
- 30 sequence-order correlation features derived from three physicochemical properties with 10 lag terms each.
- 1 sequence length feature.

The `AAC` features are:

```
AAC_A, AAC_C, AAC_D, AAC_E, AAC_F, AAC_G, AAC_H, AAC_I, AAC_K, AAC_L,
AAC_M, AAC_N, AAC_P, AAC_Q, AAC_R, AAC_S, AAC_T, AAC_V, AAC_W, AAC_Y
```

The 30 PseAAC lag features are organized into three property groups:

- Hydrophobicity: `PseAAC_hydro_lag1` to `PseAAC_hydro_lag10`.
- Polarity: `PseAAC_polar_lag1` to `PseAAC_polar_lag10`.
- Charge: `PseAAC_charge_lag1` to `PseAAC_charge_lag10`.

The sequence length feature is `Sequence_Length`.

#### Note S4. Detailed definition of CTD features

The `CTD` block contains 147 features. Seven physicochemical properties are each encoded by 21 descriptors:

- 3 composition features.
- 3 transition features.
- 15 distribution features.

The seven grouping schemes are:

- Hydrophobicity
- Normalized van der Waals volume
- Polarizability
- Secondary structure propensity
- Solvent accessibility
- Polarity
- Charge

For each property, amino acids are partitioned into three groups, and the following descriptor families are computed:

- `CTD_<property>_Comp_G1`, `CTD_<property>_Comp_G2`, `CTD_<property>_Comp_G3`
- `CTD_<property>_Trans_G1G2`, `CTD_<property>_Trans_G1G3`, `CTD_<property>_Trans_G2G3`
- `CTD_<property>_Dist_G1_First`, `CTD_<property>_Dist_G1_25%`, `CTD_<property>_Dist_G1_50%`, `CTD_<property>_Dist_G1_75%`, `CTD_<property>_Dist_G1_Last`
- `CTD_<property>_Dist_G2_First`, `CTD_<property>_Dist_G2_25%`, `CTD_<property>_Dist_G2_50%`, `CTD_<property>_Dist_G2_75%`, `CTD_<property>_Dist_G2_Last`
- `CTD_<property>_Dist_G3_First`, `CTD_<property>_Dist_G3_25%`, `CTD_<property>_Dist_G3_50%`, `CTD_<property>_Dist_G3_75%`, `CTD_<property>_Dist_G3_Last`

#### Note S5. Detailed definition of WinAAC features

To complement CTD, sxLaep further incorporates Windowed Amino Acid Composition (WinAAC) to capture local compositional variations along the protein sequence backbone. For a sequence of length $L$, the sequence is partitioned into a small number of consecutive windows defined on relative positions so that variable-length sequences can be compared. Within each window, a 20-dimensional amino acid composition vector is computed, and the window-level vectors are then concatenated to form the final WinAAC representation.

The `WinAAC` block contains 60 features and represents amino acid composition in three sequence segments:

- Segment 1: `WinAAC_Seg1_A` to `WinAAC_Seg1_Y`.
- Segment 2: `WinAAC_Seg2_A` to `WinAAC_Seg2_Y`.
- Segment 3: `WinAAC_Seg3_A` to `WinAAC_Seg3_Y`.

This design provides localized compositional information that is not fully captured by global descriptors alone and complements the grouped physicochemical patterns captured by `CTD`.

#### Note S6. Feature naming conventions

The full feature list follows these naming conventions:

- `AAC_*`: global amino acid composition
- `PseAAC_*`: sequence-order correlation descriptors
- `CTD_*_Comp_*`: grouped composition descriptors
- `CTD_*_Trans_*`: grouped transition descriptors
- `CTD_*_Dist_*`: grouped distribution descriptors
- `WinAAC_Seg*_*`: local amino acid composition in sequence segments

These feature definitions provide the basis for reproducible implementation, feature-importance interpretation, and downstream model explainability analyses.

#### Note S7. Evaluation metric formulas

We define the binary classification outcomes as true positives (`TP`), true negatives (`TN`), false positives (`FP`), and false negatives (`FN`). Let the total number of samples be

$$N = TP + TN + FP + FN$$

The evaluation metrics used in this study are defined as follows.

**Accuracy**

$$\text{Accuracy} = \frac{TP + TN}{TP + TN + FP + FN}$$

**Precision**

$$\text{Precision} = \frac{TP}{TP + FP}$$

**Recall (Sensitivity)**

$$\text{Recall} = \text{Sensitivity} = \frac{TP}{TP + FN}$$

**Specificity**

$$\text{Specificity} = \frac{TN}{TN + FP}$$

**False Negative Rate (FNR)**

$$\text{FNR} = \frac{FN}{TP + FN} = 1 - \text{Recall}$$

**F1-score**

$$\text{F1} = \frac{2 \times \text{Precision} \times \text{Recall}}{\text{Precision} + \text{Recall}}$$

**ROC-AUC**

The receiver operating characteristic area under the curve (`ROC-AUC`) is defined as the area under the ROC curve formed by plotting the true positive rate against the false positive rate across all decision thresholds:

$$\text{TPR} = \frac{TP}{TP + FN}, \qquad \text{FPR} = \frac{FP}{FP + TN}$$

and

$$\text{ROC-AUC} = \int_0^1 \text{TPR}(\text{FPR})\, d(\text{FPR})$$

**Delta recall**

For the remote-homology comparison between sxLaep and Diamond, the difference in recall is

$$\Delta\text{Recall} = \text{Recall}_{\text{sxLaep}} - \text{Recall}_{\text{Diamond}}$$

**Standard error (SE) of recall**

Standard errors were estimated using bootstrap resampling. Let $B$ denote the number of bootstrap replicates, let $\text{Recall}^{(b)}$ denote the recall estimated from the $b$-th bootstrap sample, and let $\overline{\text{Recall}}$ denote the mean recall across bootstrap replicates. The bootstrap standard error of recall is the empirical standard deviation of the bootstrap estimates:

$$\text{SE}(\text{Recall}) = \sqrt{\frac{1}{B-1}\sum_{b=1}^{B}\left(\text{Recall}^{(b)} - \overline{\text{Recall}}\right)^2}$$

**Standard error of delta recall**

For each bootstrap replicate $b$, we computed the recall difference between sxLaep and Diamond on the same resampled data:

$$\Delta\text{Recall}^{(b)} = \text{Recall}^{(b)}_{\text{sxLaep}} - \text{Recall}^{(b)}_{\text{Diamond}}$$

Let $\overline{\Delta\text{Recall}}$ denote the mean recall difference across bootstrap replicates. The bootstrap standard error of delta recall is the empirical standard deviation of these bootstrap differences:

$$\text{SE}(\Delta\text{Recall}) = \sqrt{\frac{1}{B-1}\sum_{b=1}^{B}\left(\Delta\text{Recall}^{(b)} - \overline{\Delta\text{Recall}}\right)^2}$$

##### Note S8. Hardware and software environment

All experiments were performed on a Linux server running Ubuntu 22.04.5 LTS (Jammy Jellyfish). The software environment used Python 3.11.5. All reported experiments were run under the same operating system and software environment.
